## Supporting Information for "Quantifying the impact of ecological memory on the dynamics of interacting communities"

### S1 Appendix: Methodological details for Fig 5c and S1 Fig

**Fig 5c**: Ternary plots allow representing the state of a 3-species or 3-group system by a single dot and therefore are a convenient way to display the outcome of many simulations at a time. In Fig 5c, each ternary plot shows the stable state distribution of the group relative abundances obtained for 50 different simulations, each represented by a dot of the color of the dominant group. We detail below how we computed the position of each dot in a triangle (Fig A1). Let us write $B$, $G$ and $R$ the average stable state

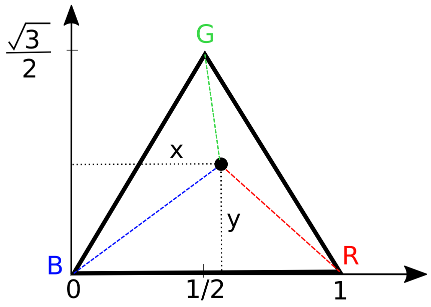


**Fig A1.** Triangle coordinates.

relative abundances of the species in the blue, green and red groups, that is $R = \frac{\sum_{i=1}^{5} R_i(end)}{\sum_i (R_i + B_i + G_i)(end)}$ (and similarly for $B$ and $G$), where $Z_i(end)$ denotes the abundance of species $i$ in group $Z$ at the end of the simulation. Let us consider an equilateral triangle in which each vertex corresponds to the complete dominance of one group of species, as shown in Fig A1. Thus, a point (dot) close to the middle of the triangle indicates a state of the system characterized by relatively even species abundances. If $B = 1$ (100%) is placed at $(x, y) = (0, 0)$ and $R = 1$ (100%) at $(1, 0)$, then $G = 1$ (100%) is at $(\frac{1}{2}, \frac{\sqrt{3}}{2})$, and any triplet $(B, R, G)$ will be at $(x, y) = \left(\frac{1}{2}(2R + G), \frac{\sqrt{3}}{2}G\right)$. These Cartesian coordinates provide a way to map any triplet of group relative abundances to a unique location on the triangle.

**S1 Fig**: Here, we randomly generated an interaction matrix $\mathbf{K}$ without predefined structure between $N = 15$ species. Specifically, we set $n = 4$ and $K_{ij} = 1 - e^{-5z}$, where $z$ is a randomly generated number from a uniform distribution between 0 and 1. We generated 10 communities, each with a random vector of growth rates generated as $b_i \sim \mathcal{N}(1, 0.0025)$, $\forall i$. We used the same interaction matrix for all 10 communities, and death rates $k_i = 2$, $\forall i$. For each community, we set the initial values for species abundances $X_i$ at one of the equilibrium points of the system (randomly chosen). To compute the dissimilarity of the community between times $t_r$ and $t_p$, we used the Bray-Curtis distance, computed as $BC(t_r, t_p) = \frac{\sum_{i=r}^{N} |X_i(t_r) - X_i(t_p)|}{\sum_{i=1}^{N} X_i(t_r) + X_i(t_p)}$.

### S2 Appendix: Numerical simulations

In the following, we describe the numerical algorithm that we used to solve fractional-order differential equation systems.

Adams methods provide commonly used numerical solutions for ordinary differential equations, involving implicit (Adams-Moulton) and explicit (Adams-Bashforth) linear multi-step schemes. We exploited in this paper the predictor-corrector method based on Adams formulae (see [79]).

Given the system equation (3) in the Methods, let us write $\mathbf{X}$ the set of all species abundances, $\boldsymbol{\mu}$ the corresponding vector of derivative orders $\mu_i$, and $\mathbf{F}$ the corresponding matrix function of all $F_i = X_i\,(b_i f_i(\{X_k\}) - k_i X_i)$. We can then rewrite the fractional order model (3) in the following matrix form:

$$\mathfrak{D}^{\boldsymbol{\mu}}\mathbf{X} = \mathbf{F}(t, \mathbf{X}), \text{ where } \mathbf{X}(t_0) = \mathbf{X}_0. \tag{i}$$

The initial value problem (i) is equivalent to the Volterra integral equation [80]:

$$\mathbf{X}(t) = \mathbf{X}_0 + \frac{1}{\Gamma(\boldsymbol{\mu})}\int_{t_0}^{t} (t-\tau)^{\boldsymbol{\mu}-1}\mathbf{F}(\tau, \mathbf{X}(\tau))d\tau. \tag{ii}$$

We solved Eq. (ii) using a product integration technique, in which we replaced the function $\mathbf{F}(\tau, \mathbf{X}(\tau))$ with piece-wise interpolating polynomials. For the grid nodes $t_j$ $(j = 0, ..., m)$ with constant step size $h$ $(t_j = t_0 + jh)$, we write $\mathbf{F}_j = \mathbf{F}(t_j, \mathbf{X}_j)$ where $\mathbf{X}_j$ is the numerical approximation to $\mathbf{X}(t_j)$. The product rectangle rule [80] gives an explicit estimation of Eq. (ii) as a predictor:

$$\begin{aligned} \mathbf{X}_m &= \mathbf{X}_0 + h^{\boldsymbol{\mu}} \textstyle\sum_{j=0}^{m-1} \mathbf{b}_{m-j-1}\mathbf{F}_j, \\ \mathbf{b}_{m-j-1} &= \tfrac{(m-j)^{\boldsymbol{\mu}} - (m-j-1)^{\boldsymbol{\mu}}}{\Gamma(\boldsymbol{\mu}+1)}, \end{aligned} \tag{iii}$$

and the product trapezoidal rule [80] provides an implicit estimation of Eq. (ii) as a corrector:

$$\begin{aligned} \mathbf{X}_m &= \mathbf{X}_0 + h^{\boldsymbol{\mu}}\mathbf{c}_m\mathbf{F}_0 + h^{\boldsymbol{\mu}} \textstyle\sum_{j=1}^{m} \mathbf{d}_{m-j}\mathbf{F}_j, \\ \mathbf{c}_m &= \tfrac{(m-1)^{\boldsymbol{\mu}+1} - m^{\boldsymbol{\mu}}(m-\boldsymbol{\mu}-1)}{\Gamma(\boldsymbol{\mu}+2)}, \\ \mathbf{d}_{m-j} &= \begin{cases} \tfrac{1}{\Gamma(\boldsymbol{\mu}+2)}, & \text{if } m-j=0, \\ \tfrac{(m-j-1)^{\boldsymbol{\mu}+1} - 2(m-j)^{\boldsymbol{\mu}+1} + (m-j+1)^{\boldsymbol{\mu}+1}}{\Gamma(\boldsymbol{\mu}+2)}, & \text{if } m-j = 1, 2, ... \end{cases} \end{aligned} \tag{iv}$$

The last term of the sum in the corrector equation (iv), $\mathbf{F}(t_m, \mathbf{X}_m)$, is obtained by an approximation of $\mathbf{X}_m$ in the predictor equation (iii). This method is called FracPECE (Fractional Predict Evaluate Correct Evaluate) [80]. Because its standard implementation was not sufficient considering the stiffness of the equation, we improved its accuracy via an advanced convolution quadrature and Fast Fourier Transform [79], and via multiple applications of the corrector step [81] when required. Specifically, we used several corrector iterations when the difference between two consecutive iterations was larger than the desired tolerance of $10^{-6}$.

Note that since the model with fractional order derivatives (3) includes the standard model (1) as a particular case (namely, for integer derivative order), the numerical approximations (iii) and (iv) are also solutions to equation (1). The explicit solution (iii)–or an assessment of the implicit solution (iv)–shows how memory influences the fundamental system dynamics through the dependence on $\mu$.

**Table S1. Exact model specifications for the 2, 3, and 15-species Gonze model (equation 1 in Methods).**

| **Figure** | Gonze model (1) | | | | | | | | | | | |
|---|---|---|---|---|---|---|---|---|---|---|---|---|
| Commensurate | $X_0$ | | | $K_{ij}$ | $n$ | $k_i$ | $\mu$ | | | $b$ | | |
| | B | R | G | $\forall i \neq j$ | | $\forall i$ | B=R=G | | | B | R | G |
| **2a** | 0.99 | 0.01 | 0.01 | 0.1 | 2 | 1 | 1 & 0.9 | | | Pulse1 | 0.95 | Pulse1 |
| **2b** | | | | | | | 1 & 0.9 | | | Pulse2 | | Pulse2 |
| **3a** | | | | | | | 1 & 0.96 & 0.9 | | | Pulse3 | | 1.05 |
| **3b** | | | | | | | 1 & 0.96 & 0.9 | | | Periodic | | 1.05 |
| **4b-c** | 1/3 | 1/3 | 1/3 | 0.1 | 2 | 1 | 1<br>0.9157959<br>0.9157954<br>0.9157952<br>0.9<br>0.7<br>0.6 | | | Stochastic | | |
| Incommensurate | | | | | | | B | R | G | | | |
| **S1** | Equilibrium points | | | Random interactions | 4 | 2 | No Specified Groups $\mu_i$, $\forall i = 1$ (or 0.7) | | | $\mathcal{N}(1, 0.0025)$ with a pulse | | |
| **5** | $Uniform(0, 0.1)$ | | | Predefined interactions | 2 | 1 | 1<br>0.6<br>0.851841<br>0.851840<br>0.8 | 1 | 1 | $\mathcal{N}(1, 0.01)$ | | |
| **S2** | | | | | | | permutation of 1, 0.9, 0.6 | | | | | |
| **S3a** | 0.99 | 0.01 | 0.01 | 0.1 | 2 | 1 | 1<br>1<br>1 | 1<br>1<br>1 | 1<br>0.90895<br>0.90893 | Pulse4 | 0.95 | 1.05 |
| **S3b** | 1/3 | 1/3 | 1/3 | 0.1 | 2 | 1 | 1<br>1<br>1 | 1<br>1<br>0.9 | 1<br>0.8<br>1 | Stochastic | | |
| **S4a**<br>**S4b** & **6** | [0.005,0.05] | 0.1&0.3 | 1&0.1 | 0.1 | 2 | 1 | 1<br>0.6 | 1<br>0.6 | 1<br>1 | 4 | 0.95 | 1.05 |

| **2-Species** | $X_0$ | | $K_{ij}$ | $n$ | $k_i$ | $\mu$ | | $b$ | | Convergence interval |
|---|---|---|---|---|---|---|---|---|---|---|
| | B | R | $\forall i \neq j$ | | $\forall i$ | B | R | B | R | |
| **S5a & b** | 0.8 & 0.9 | 0.2 & 0.15 | 0.1 | 2 | 1 | [0.6,1] | [0.6,1] | 1 | 2 | - |
| **S10** | 0.9 | 0.2 | | | | [0.84,1] | [0.84,1] | | | 5e-3 |
| **S6** | Equilibrium points | | | | | [0.9,1] | [0.9,1] | Pulse5 | | - |
| **S8** | | | | | | | | | | 0.02 & 1e-6 |

Pulse1: $b_B(t) = 0.5$ and $b_G(t) = 2$ if $20 \leq t < 60$, otherwise $b_B(t) = 1$ and $b_G(t) = 1.05$.
Pulse2: $b_B(t) = 0.5$ and $b_G(t) = 2.2$ if $20 \leq t < 60$, otherwise $b_B(t) = 1$ and $b_G(t) = 1.05$.
Pulse3: $b_B(t) = 0.2$ if $60 \leq t < 100$, $b_B(t) = 4.5$ if $200 \leq t < 330$, otherwise $b_B(t) = 1$.
Periodic: $b_B(t) = 1$ if $20(2m - 2) \leq t < 20(2m - 1)$, $b_B(t) = 0.2$ if $20(4m - 3) \leq t < 20(4m - 2)$, $b_B(t) = 4.5$ if $20(4m - 1) \leq t < 20(4m)$ where $m \in \mathbb{N}$.
Stochastic: The growth rates of these panels are generated by mean-reverting the Ornstein-Uhlenbeck Process described by the stochastic equation $db_t = \theta(\phi - b_t)dt + \sigma dW_t$.
Random interactions: $K_{ij} = 1 - e^{-5z}$, where z is randomly generated from a uniform distribution between 0 and 1.
Predefined interactions: $K_{ij} \sim 1 + \mathcal{N}(0, 0.01)$ for species $i$ and $j$ in the same group (intra-group interactions $K_{BB}$, $K_{RR}$, $K_{GG}$), and $K_{ij} \sim 0.5 + \mathcal{N}(0, 0.01)$ for species $i$ and $j$ in different groups (inter-group interactions).
Pulse4: $b_B(t) = 0.2$ if $60 \leq t < 100$, $b_B(t) = 4.5$ if $400 \leq t < 530$, otherwise $b_B(t) = 1$.
Pulse5: $b_R(t) = 2$ and $b_B(t) = b_B(t) + p$ if $50 \leq t < 100$, otherwise $b_B(t) = 1$, where $p$ is a positive value.

**Table S2. Exact model specifications for the 2-species model given by equation 2 in Methods, and for the logistic growth curve in Fig 7.**

| **Logistic growth curve** | | $X_0$ | | $\kappa$ | | $\mu$ | | $b$ | | | |
|---|---|---|---|---|---|---|---|---|---|---|---|
| **Fig 7** | | 0.1 | | 1 | | [0.6:1] | | 1 | | | |
| **Model (2)** | $X_0$ | | $K_{ij}$ | | | | $\mu$ | | $b$ | | Convergence |
| **Figure** | BU | BT | BUBU | BUBT | BTBU | BTBT | BU | BT | BU | BT | interval |
| **S5c & d** | 0.3 & 0.28 | 0.36 | -0.9059 | -0.9377 | -0.9720 | -0.9597 | [0.6,1] | [0.6,1] | 0.599 | 0.626 | - |
| **S11** | 0.3 | 0.37 | | | | | [0.84,1] | [0.84,1] | | | 5e-3 |
| **S7** | Equilibrium points | | | | | | [0.9,1] | [0.9,1] | Pulse6 | | - |
| **S9** | | | | | | | [0.9,1] | [0.9,1] | | | 0.02 & 7e-4 |
| | CH | ER | CHCH | CHER | ERCH | ERER | CH | ER | CH | ER | Convergence interval |
| **S5e** | 0.4 | 0.2 | -1.2420 | -0.5077 | 1.1905 | -1.3219 | [0.6,1] | [0.6,1] | 0.468 | 0.151 | - |
| **S12** | | | | | | | [0.9,1] | [0.9,1] | | | 1e-4 |
| | BT | CH | BTBT | BTCH | CHBT | CHCH | BT | CH | BT | CH | Convergence interval |
| **S5f** | 0.158 | 0.8 | -0.9597 | -0.0727 | -0.5906 | -1.2420 | [0.6,1] | [0.6,1] | 0.626 | 0.468 | - |
| **S13** | | | | | | | [0.9,1] | [0.9,1] | | | 1e-3 |

Pulse6: $b_{BT}(t) = 0.626$ and $b_{BU}(t) = b_{BU}(t) + p$ if $50 \leq t < 80$, otherwise $b_{BU}(t) = 0.599$, where $p$ is a positive value.

Logistic growth curve: a well-known model of biological population dynamics, which describes simple exponential growth. $\frac{dX}{dt} = b\left(1 - \frac{X}{\kappa}\right)X$, where $X$ represents the population size, $b$ the growth rate, and $\kappa$ the carrying capacity.

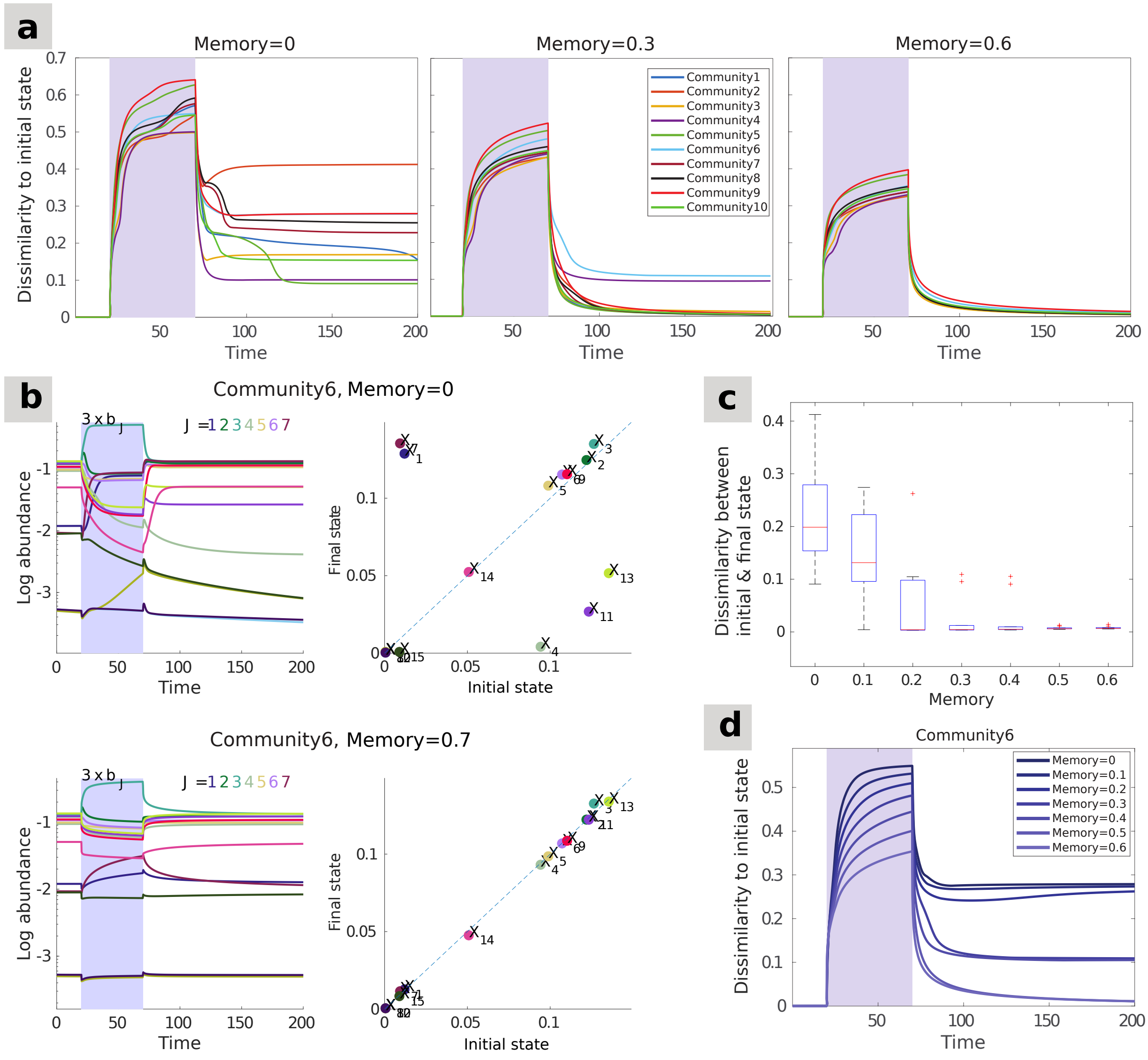


**Fig S1. Memory effects preserve the stable state in randomly structured communities.** We simulated ten communities of 15 species each with random interaction matrices (see S1 Appendix for details). We applied a similar level of commensurate memory to all ten communities. Every community is initially in a stable state of the system, and a perturbation is imposed by multiplying the growth rates of half of the species $(b_1, ..., b_7)$ by 3. The simulation is stopped when the system is close to its new stable state. Although only the effect of commensurate memory is illustrated here, the same outcome can be achieved using incommensurate memory. **(a)** Dissimilarity (Bray-Curtis) to the initial stable state through time for all ten communities, for three different memory strengths. The stronger the memory, the more constrained the community trajectories are, and the more likely they are to revert to their initial stable state eventually. **(b)** Time series for one randomly chosen community, community 6. The pulse perturbations lead the community to an alternative stable state in the absence of memory (top), while adding memory effects allows recovering the original state (bottom). **(c)** Community dissimilarity (Bray-Curtis) between the start and the end of the simulation for all ten communities and different memory levels. Without memory, the pulse perturbation changes the abundances of some of the species and leads to an alternative stable state (*i.e.*, non-zero dissimilarity between start and end). In contrast, all communities recover their pre-perturbation stable state in the presence of memory (*i.e.*, zero dissimilarity). **(d)** Dissimilarity to the initial stable state through time in community 6, for different memory strengths.

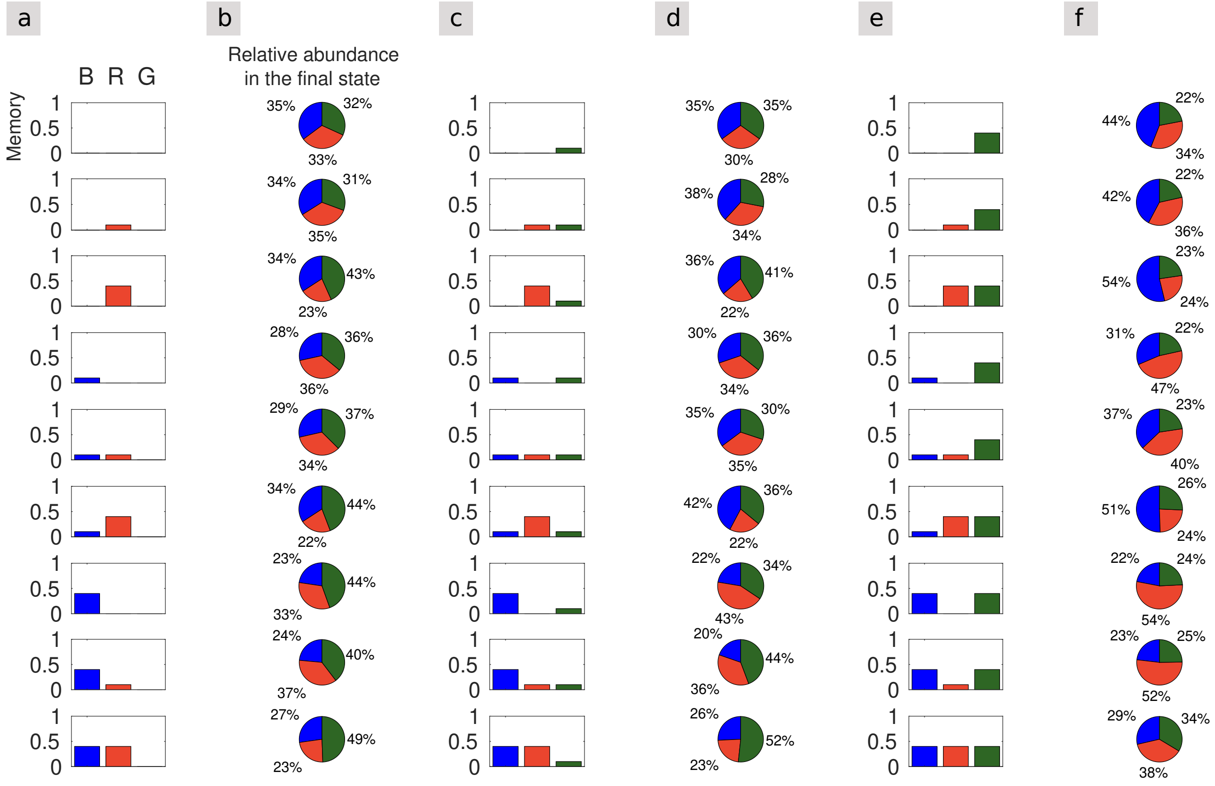


**Fig S2. Memory in a group of species decreases their relative abundance.** Same as Fig 5c, for varying incommensurate memory values in the three species groups. For each of the 27 memory configurations represented in columns a, c and e (*i.e.*, varying memory strengths for species groups blue, red and green), the same model as in Fig 5 was simulated 50 times with random initial conditions. Columns b, d and e show the corresponding outcomes, represented as the mean relative abundance achieved by each species group in the final state across the 50 simulations. The columns are ordered by increasing strength of memory in the green species group (from left to right), and the rows by increasing strength of memory in the blue species group (from top to bottom). For each of the three groups, increasing the strength of their memory leads to decreasing their relative abundance in the final state.

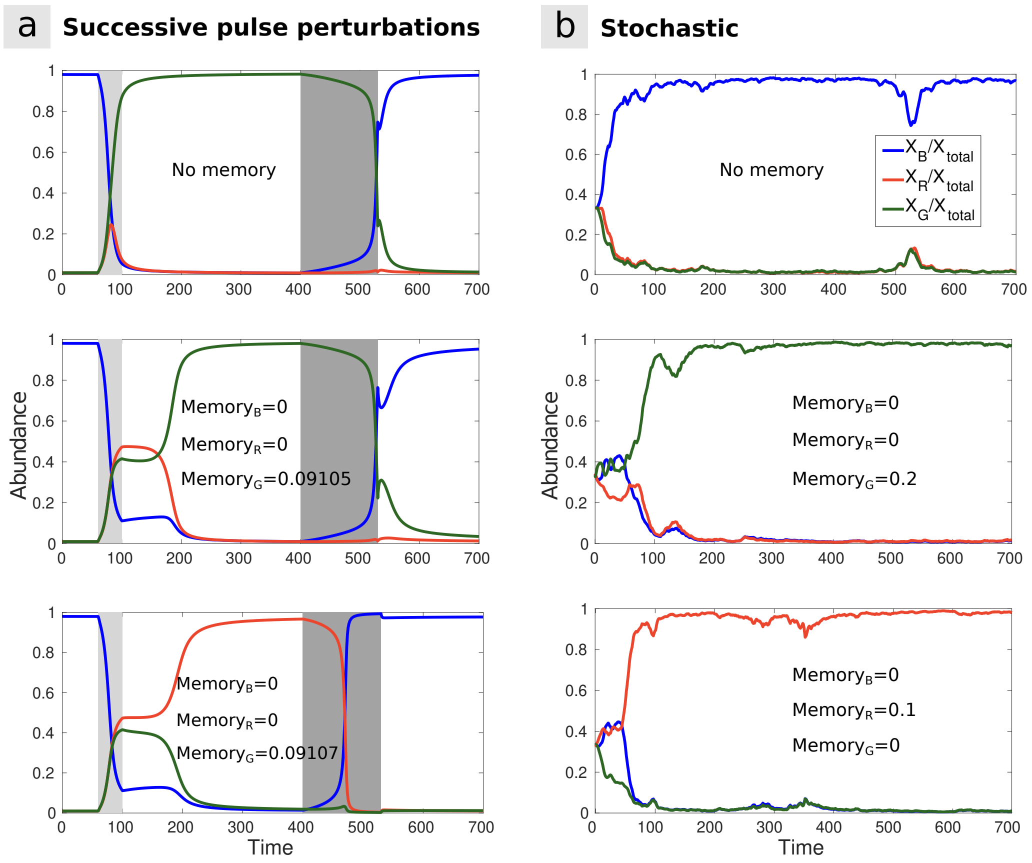


**Fig S3. Impact of incommensurate memory in the presence of perturbation.** **(a)** Same as Fig 3a-b but with incommensurate memory: memory is applied to the green species with increasing strength from top to bottom, while the blue and red species remain memoryless. Around a particular memory strength (0.09106), the system changes behavior: when memory in the green species is between 0 and 0.09105, the first perturbation leads to the green species achieving dominance, whereas between 0.09107 and 1, it leads to the red species achieving dominance. **(b)** Same as Fig 4 but with incommensurate memory. In the absence of memory, the blue species is dominant. However, when imposing sufficient memory on the green or red species, they respectively become dominant in the stable state.

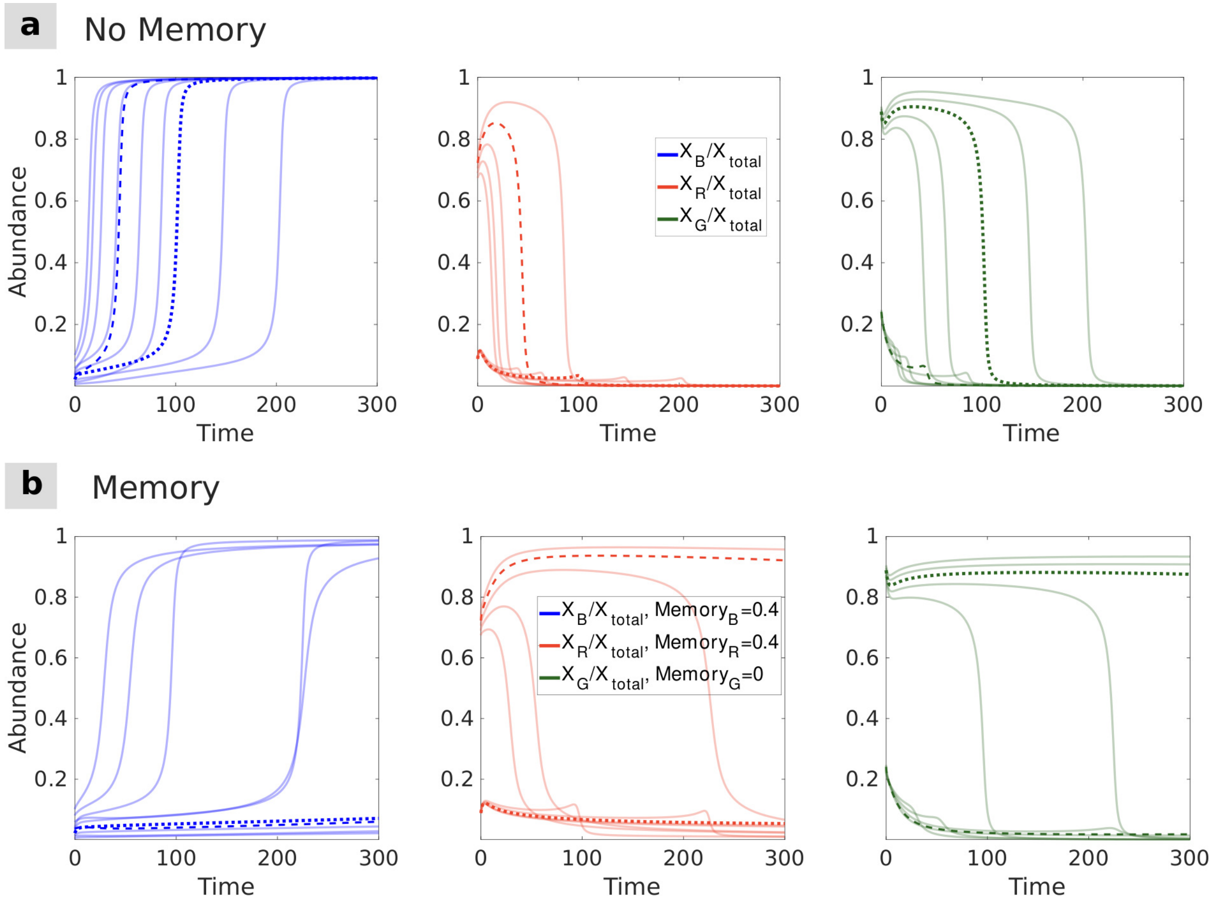


**Fig S4. Memory can induce long transient dynamics even in the absence of multistability.** Three-species community model converging to a single stable state irrespective of initial conditions, but close enough to the tristable region in the model's parameter space that the introduction of memory induces alternative long transient states (see Fig 6). In each row, the three panels show the relative abundances of the blue, red and green species along time for the same 10 simulations starting from 10 different initial conditions (*i.e.*, sets of initial abundances). **(a)** In the absence of memory, the dynamics quickly converges to the stable state irrespective of initial conditions. **(b)** In the presence of memory, the dynamics may, depending on the initial conditions, remain stuck in alternative transient states far from the stable state (although it would eventually converge to the stable state should the simulation be run long enough). The dashed and dotted lines indicate the initial abundance thresholds that separate from each other the stable state and the two alternative transient regimes, each corresponding to the dominance of a different species. The same initial conditions are also indicated by dashed and dotted lines in (a)

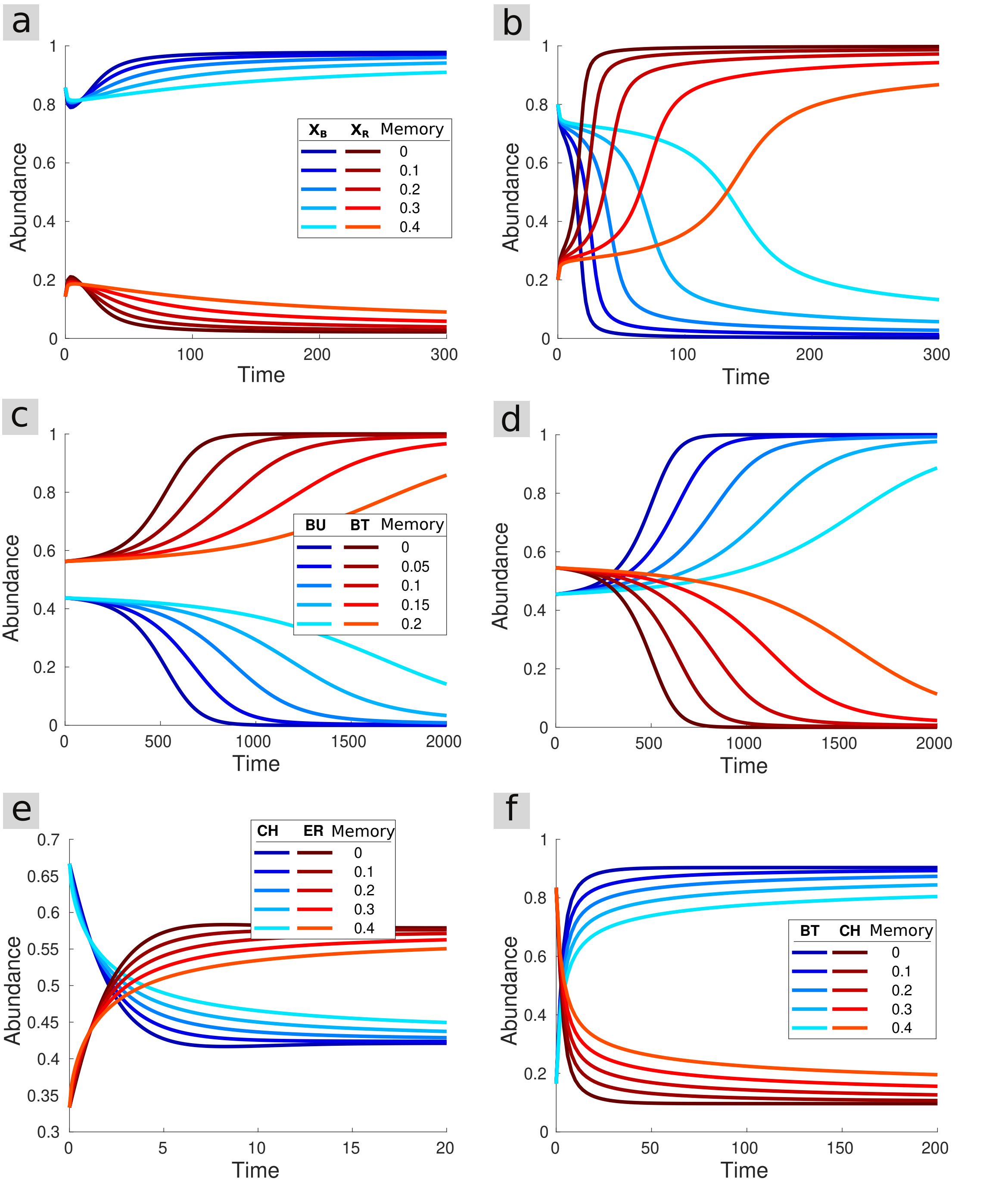


**Fig S5. Memory effects on convergence time for different two-species community types.** Impact of memory on the dynamics of the different communities considered in the "Empirically parameterized model" section of the results, with a range of initial abundances for each panel. **(a-b)** Two-species version of the multistable (here bistable) community model from Gonze model (described by equation (1) in the Methods). (a) and (b) show a set of initial abundances leading to one or the other stable state (blue- or red-dominated). **(c-d)** Bistable community composed of *Bacteroides uniformis* (BU) and *Bacteroides thetaiotaomicron* (BT) (model described by equation (2) in the Methods, as for panels (e) and (f)). (c) and (d) show a set of initial abundances leading to one or the other stable state (BT- or BU-dominated, respectively). **(e)** Monostable community exhibiting coexistence between *Eubacterium rectale* (ER) and *Clostridium hiranonis* (CH) in the stable state. **(f)** Monostable community exhibiting dominance of *Bacteroides thetaiotaomicron* (BT) over *Clostridium hiranonis* (CH) in the stable state. The corresponding parameter values are detailed in Table in S2 Table.

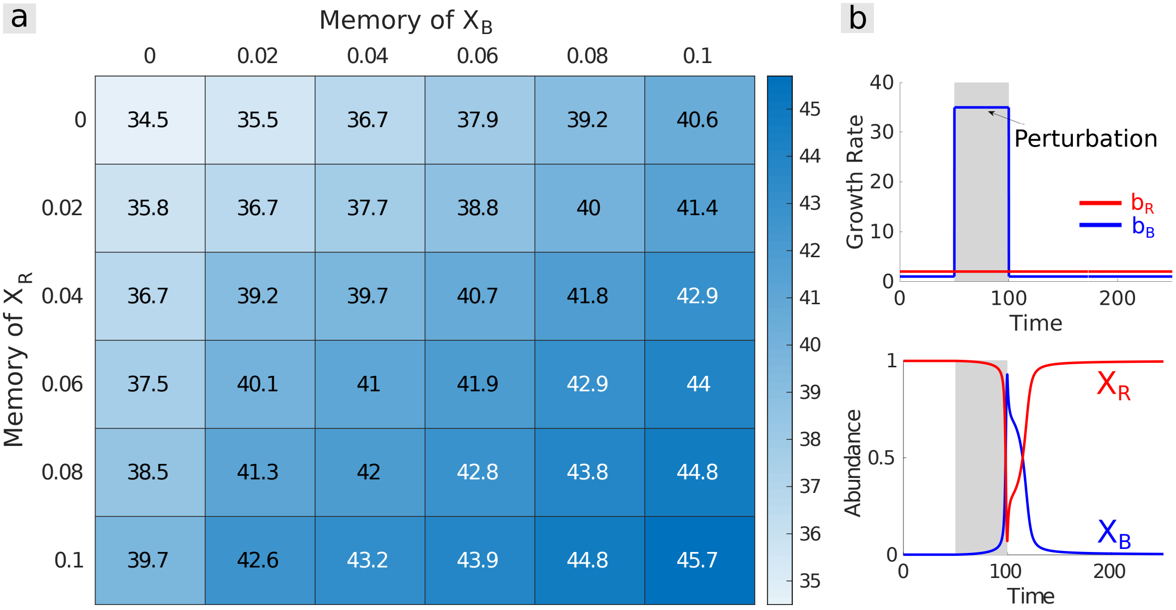


**Fig S6. Impact of memory on resistance to a pulse perturbation in the two-species version of Gonze multistable model. (a)** Both color and matrix entries indicate the strongest pulse perturbation for which the community recovers to its initial stable state, as a function of memory strength in the blue and red species (abundances $X_B$ and $X_R$, respectively). Increasing memory in either species increases system resistance in a similar way. **(b)** For each matrix entry in (a), a pulse perturbation is applied to the growth rate of the blue species (top panel), which temporarily displaces the community away from its original stable state dominated by the red species (bottom panel). The value taken by the blue species growth rate during the pulse determines the intensity of the perturbation.

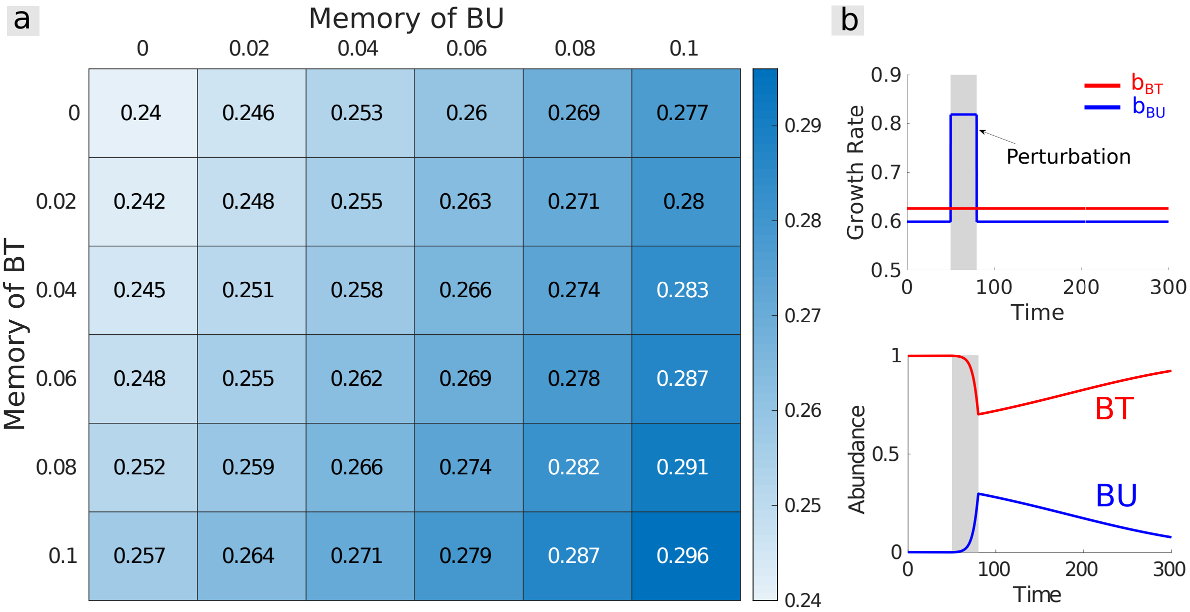


**Fig S7. Impact of memory on resistance to a pulse perturbation in a two-species community exhibiting bistability between dominance of *Bacteroides uniformis* (BU) and *Bacteroides thetaiotaomicron* (BT). (a)** Both color and matrix entries indicate the strongest pulse perturbation for which the community recovers to its initial stable state, as a function of memory strength in *BU* and *BT*. As in S6 Fig, increasing memory in either species increases system resistance in a similar way. **(b)** For each matrix entry in (a), a pulse perturbation is applied to the growth rate of BU (top panel), which temporarily displaces the community away from its original stable state dominated by BT (bottom panel). The value taken by BU growth rate during the pulse determines the intensity of the perturbation.

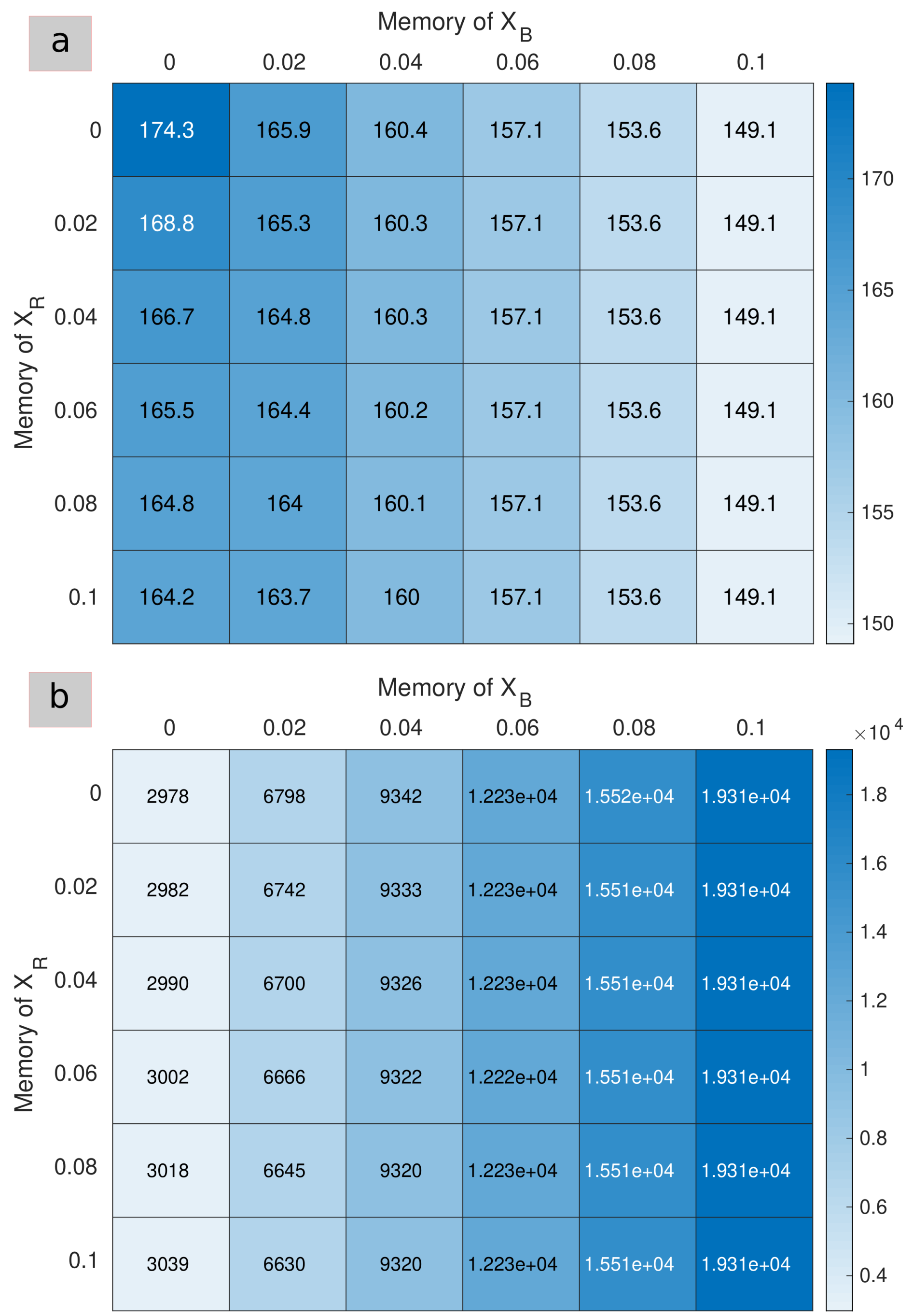


**Fig S8. Impact of memory on resilience after a pulse perturbation in the two-species version of Gonze multistable model.** Both color and matrix entries indicate the recovery time after a pulse perturbation starting from the red-dominated stable state as a function of memory strength in the blue and red species (abundances $X_B$ and $X_R$, respectively)). **(a)** Recovery times are first measured using a loose convergence interval of 0.02 on the distance of the community to its initial stable state, thus capturing the early stages of the recovery (see Resistance and resilience metrics section in Methods). Recovery time decreases with increasing memory, i.e., memory effects increase resilience. **(b)** Recovery times are then measured using a much tighter convergence interval of $1e-6$, thus capturing the later stages of the recovery. Recovery time now decreases with increasing memory, i.e., memory effects decrease resilience. Hence, the effect of memory on resilience depends on the time scale considered: memory hastens the recovery at first (a) but slows it down further in time (b). In both (a) and (b), memory in the species that is less abundant in the stable state, the blue species, has a much stronger influence on the dynamics.

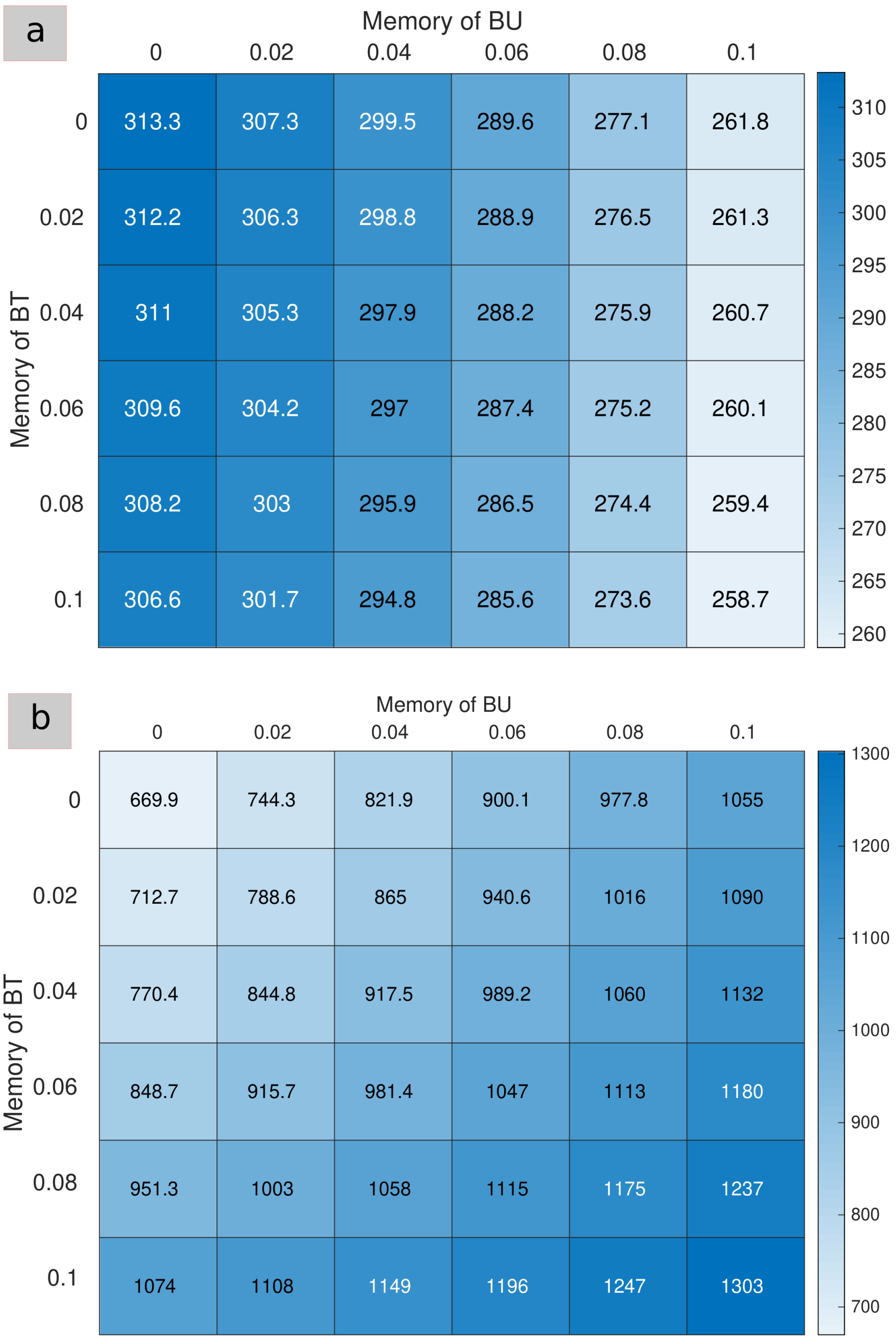


**Fig S9. Impact of memory on resilience after a pulse perturbation in a two-species community exhibiting bistability between dominance of *Bacteroides uniformis* (BU) and *Bacteroides thetaiotaomicron* (BT).** Both color and matrix entries indicate the recovery time after a pulse perturbation starting from the BT-dominated stable state, as a function of memory strength in $BU$ and $BT$. **(a)** As in S8 Fig, recovery times are first measured using a loose convergence interval of 0.02 on the distance of the community to its initial stable state, thus capturing the early stages of the recovery. Recovery time decreases with increasing memory, i.e., memory effects increase resilience. **(b)** Recovery times are then measured using a much tighter convergence interval of $7e-4$, thus capturing the later stages of the recovery. Recovery time now decreases with increasing memory, i.e., memory effects decrease resilience. Hence, as for the two-species Gonze model in S8 Fig, the effect of memory on resilience depends on the time scale considered: memory hastens the recovery at first (a) but slows it down further in time (b). As in S8 Fig, memory in the species that is less abundant in the stable state, BU, has a much stronger influence on the dynamics in (a). In (b), however, the effect of both species is of similar magnitude.

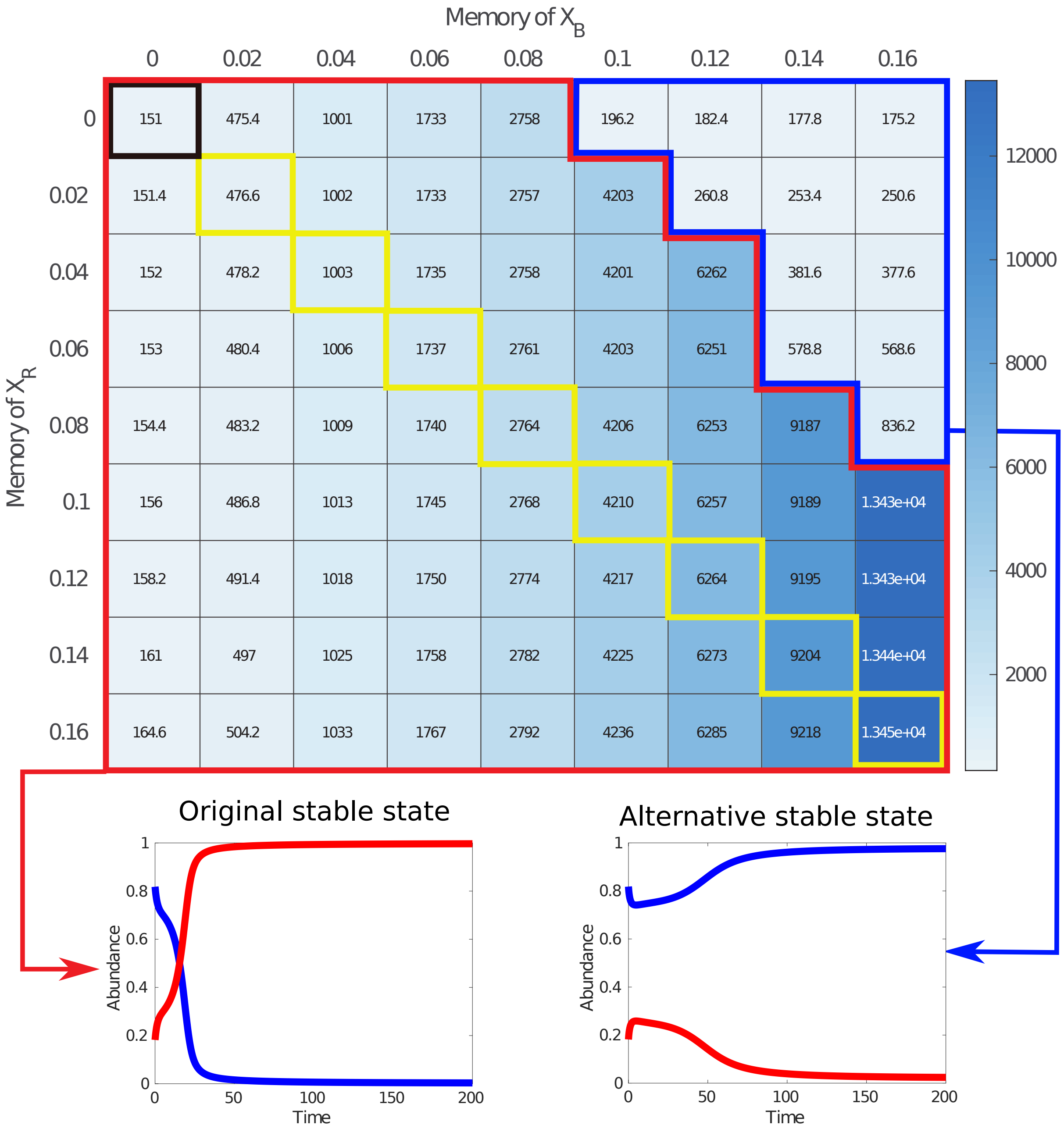


**Fig S10. Impact of memory on convergence time in the two-species version of Gonze multistable model.** Both color and matrix entries indicate the convergence time to stable state in the absence of perturbation as a function of memory strength in the blues and red species (abundances $X_B$ and $X_R$, respectively). The upper-left cell corresponds the memoryless case (thick black border). Diagonal cells correspond to commensurate memory (yellow border). Cells with a red border correspond to communities that converge to the same stable state (dominated by the red species) as the community without memory, whereas cells with a blue border correspond to communities that converge to the alternative stable state (dominated by the blue species). In the red region of the matrix, increasing memory in either species increases the convergence time to the stable state (i.e., slows down the convergence). In the blue region, increasing memory in $X_R$ also increases the convergence time, but increasing memory in $X_B$ has little effect, and actually reduces the convergence time slightly. In both regions of the matrix, increasing memory in the species that is less abundant in the stable state has a stronger influence on the dynamics.

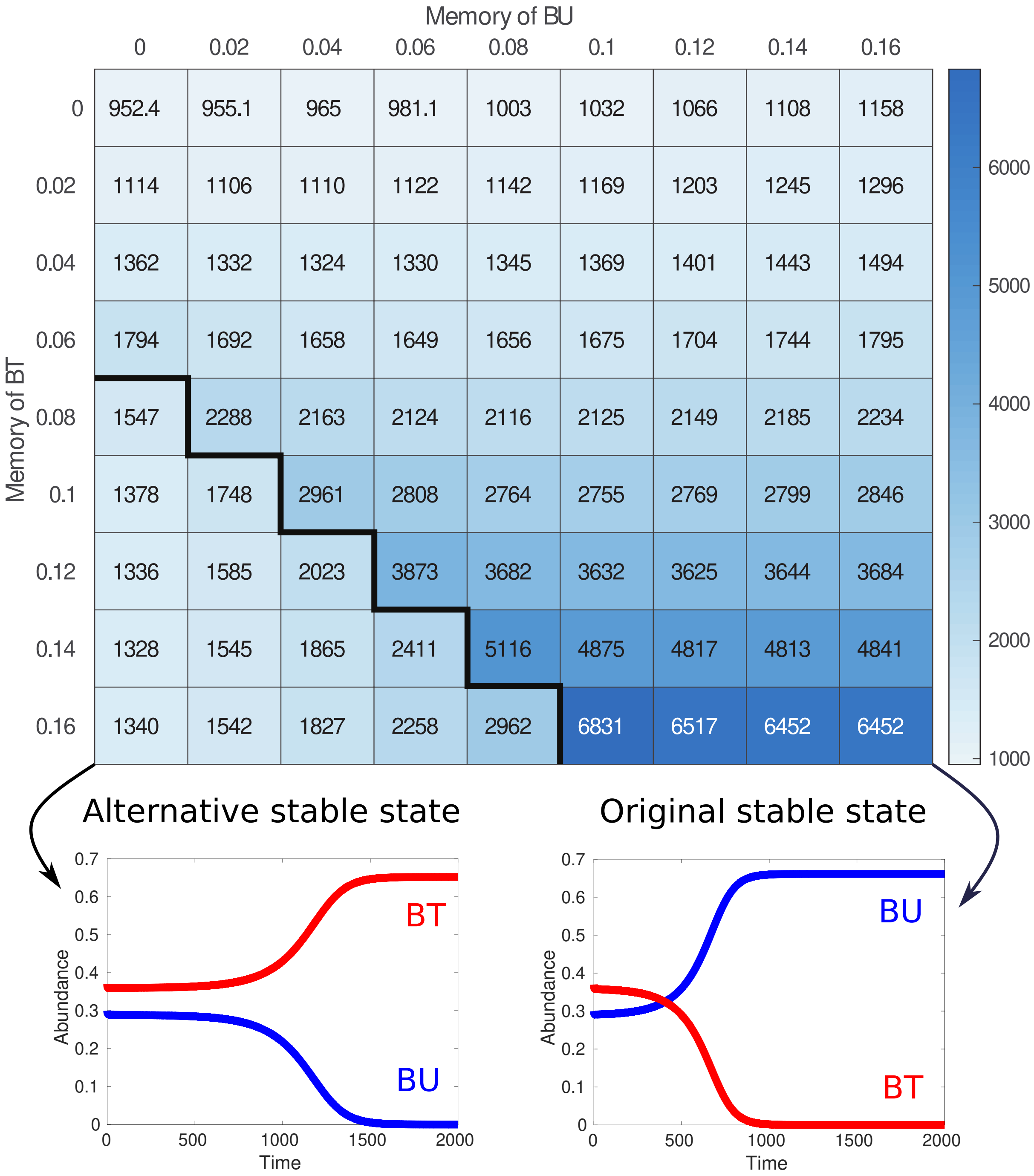


**Fig S11. Impact of memory on convergence time in a two-species community exhibiting bistability between dominance of *Bacteroides uniformis* (BU) and *Bacteroides thetaiotaomicron* (BT).** Both color and matrix entries indicate the convergence time to stable state as a function of memory strength in $X_B$ and $X_R$. The upper-left cell corresponds to the memoryless case. Diagonal cells correspond to commensurate memory. Cells above the thick black border correspond to communities that converge to the same stable state (dominated by BU) as the community without memory, whereas cells below this border correspond communities that converge to the alternative stable state (dominated by BT). In the region of the matrix above the state transition, increasing memory in either species increases the convergence time to the stable state (i.e., slows down the convergence). In the region of the matrix below the state transition, increasing memory in BU also increases the convergence time, but increasing memory in BT has little effect, and actually reduces the convergence time near the transition. In both regions of the matrix, increasing memory in the species that is less abundant in the stable state has a stronger influence on the dynamics.

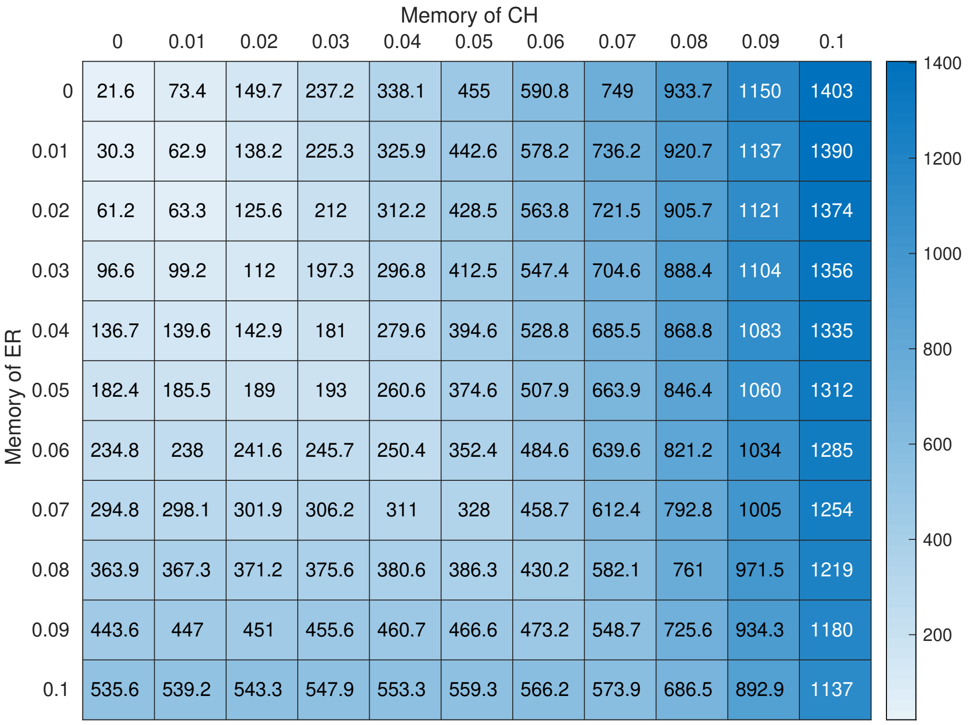


**Fig S12. Impact of memory on convergence time in a two-species community exhibiting stable coexistence between *Eubacterium rectale* (ER) and *Clostridium hiranonis* (CH).** Both color and matrix entries indicate the convergence time to stable state as a function of memory strength in ER and CH. The upper-left cell is the memoryless case, and the diagonal cells correspond to commensurate memory. Increasing memory in either species monotonically slows down the speed of convergence to the stable state, but memory in CH, the less abundant species in the stable state, has a much stronger influence.

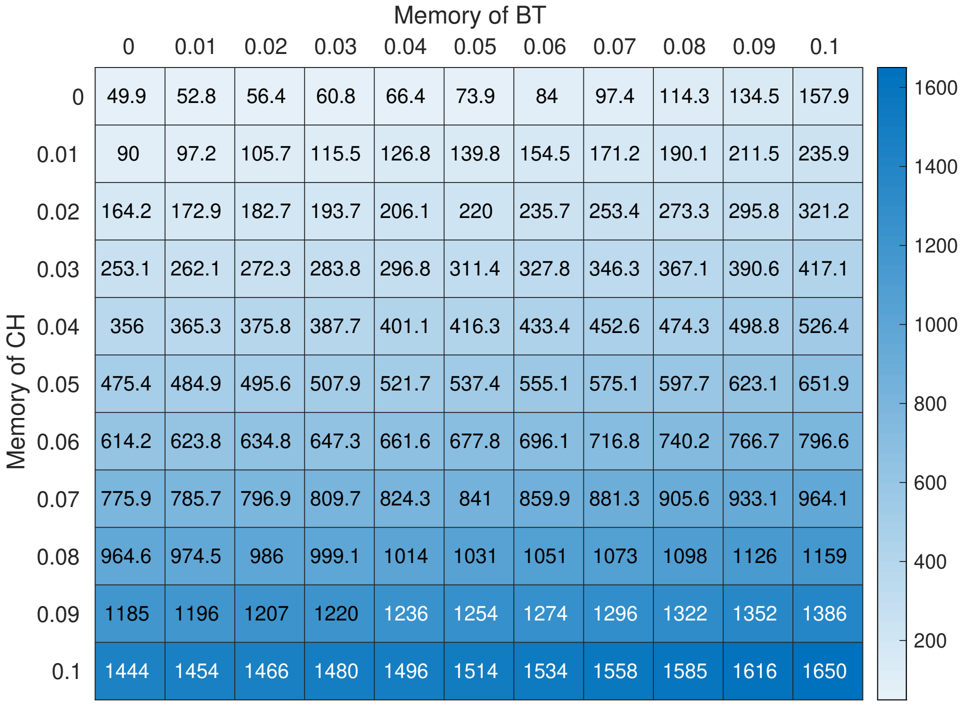


**Fig S13. Impact of memory on convergence time in a two-species community exhibiting stable dominance of *Clostridium hiranonis* (CH) by *Bacteroides thetaiotaomicron* (BT).** Both color and matrix entries indicate the convergence time to stable state as a function of memory strength in BT and CH. The upper-left cell is the memoryless case, and the diagonal cells correspond to commensurate memory. Increasing memory in either species monotonically slows down the speed of convergence to the stable state, but memory in CH, the less abundant species in the stable state, has a much stronger influence.
